# Polynuclear ruthenium complexes are effective antibiotics against *Pseudomonas aeruginosa*

**DOI:** 10.1101/2022.08.11.503708

**Authors:** Brent S. Weber, Lindsey A. Carfrae, Joshua J. Woods, Kristina Klobucar, Nicholas P. Bigham, Craig R. MacNair, Tracy L. Raivio, Justin J. Wilson, Eric D. Brown

## Abstract

There is an urgent need to develop new antibiotics for the treatment of infections caused by drug-resistant Gram-negative bacteria. In particular, new and diverse chemical classes of antibiotics are needed, as most antibiotics in clinical development are derivatives of existing drugs. Despite a history of use as antimicrobials, metals and metal-based compounds have largely been overlooked as a source of new chemical matter for antibacterial drug discovery. In this work, we identify several ruthenium complexes, ruthenium red, Ru265, and Ru360’, that possess potent antibacterial activity against both laboratory and clinical isolates of *Pseudomonas aeruginosa*. Suppressors with increased resistance were sequenced and found to contain mutations in the mechanosensitive ion channel *mscS-1* or the *colRS* two component system. The antibacterial activity of these compounds translated *in vivo* to *Galleria mellonella* larvae and mouse infection models. Finally, we identify strong synergy between these compounds and the antibiotic rifampicin, with a dose-sparing combination therapy showing efficacy in both infection models. Our findings provide clear evidence that these ruthenium complexes are effective antibacterial compounds against a critical priority pathogen and show promise for the development of future therapeutics.

## Introduction

The antimicrobial resistance crisis is a major threat to global public health, and our ability to treat serious bacterial infections remains tenuous. The pipeline of new drugs, particularly for critical priority Gram-negative pathogens, is inadequate to meet the realities facing modern healthcare (1, 2). For numerous reasons, including economic and safety concerns, antibiotic development continues to experience challenges in bringing new drugs to market (3). While most new antibiotics under development for the treatment of Gram-negative infections are derivatives of existing drug classes (1), there has been increasing interest in less conventional therapeutics (4), including anti-virulence compounds, host-directed therapeutics, and drug repurposing strategies (5–8). Despite substantial efforts, however, innovations in this space have yet to translate to successful clinical approvals of novel drugs (9).

Metals have played an important role in medicine, including their use as antimicrobial agents. One of the first synthetic antibiotics, Salvarsan, was an arsenic-based compound used to treat syphilis (10), and dressings embedded with silver formulations are used as therapeutics for burn wounds (11, 12). While some metal-based compounds have found success in other clinical applications, such as platinum metals for cancer therapies (13), there are no metal-based systemic therapies for bacterial infections. This is likely due to the historical success of conventional antibacterial therapies and the perceived toxicity concerns with metal compounds. However, with the rise of antibacterial resistance, metal-based antibiotics are drawing increased interest (14, 15). Indeed, recent work has emphasized that metal compounds are rich sources of both antibacterial and antifungal agents (16).

Recently, we showed that the metal complex ruthenium red (RuRed) possesses antibacterial activity against the Gram-negative bacterial pathogen *Klebsiella pneumoniae* under certain growth conditions, which translated to efficacy in an *in vivo* rat infection model (17). RuRed and related ruthenium complexes are primarily recognized as inhibitors of the mitochondrial calcium uniporter (MCU) (18, 19). Mitochondrial Ca^2+^ plays an important role in diverse cellular processes, and uptake of Ca^2+^ into mitochondria is mediated by the MCU ion channel (20–22). Several disease states are associated with dysregulated mitochondrial Ca^2+^ influx, including cystic fibrosis (CF), which has led to interest in therapeutics and chemical probes targeting the MCU (23, 24). In this work, we show that RuRed and other similar ruthenium complexes have potent antibacterial activity against laboratory and clinical isolates of *Pseudomonas aeruginosa*. We find that *P. aeruginosa* shows unique susceptibility to these compounds and identify mutations that decrease RuRed susceptibility. Finally, we show that these ruthenium complexes are effective as therapeutics in several *in vivo* infection models and synergize with the Gram-positive antibiotic rifampicin in dose-sparing combination treatments. Our work provides a foundation for further exploration of these ruthenium complexes as therapies to treat *P. aeruginosa* infections.

## Results

### Ruthenium red has potent antibacterial activity against laboratory and clinical isolates of *P. aeruginosa*

Previously, we found that RuRed had antibacterial activity against *K. pneumoniae* in the presence of human serum (17). In that work, tests in conventional lysogeny broth (LB) growth medium revealed RuRed did not inhibit growth of wild-type *K. pneumoniae* at concentrations up to 256 µg/mL, though we identified genetic mutants that were susceptible under those growth conditions (17). Here, we tested the activity of RuRed against two common laboratory strains of *P. aeruginosa*, PAO1 and PA14 (Fig. 1A). Surprisingly, RuRed inhibited the growth of both strains in standard antibiotic testing medium Mueller-Hinton broth (MHB), with a minimum inhibitory concentration (MIC) of 2 µg/mL (Fig. 1B). Like our previous work, growth in serum enhanced the activity of RuRed against *P. aeruginosa* PAO1 (Fig. S1) and tests in LB showed a 4-fold decrease in efficacy against PAO1 (Table S1). Interestingly, we detected an MIC for RuRed against *K. pneumoniae* in MHB (64 µg/mL), but not LB (>256 µg/mL), suggesting a difference in efficacy between LB and MHB (Table S1).

**Fig 1.**
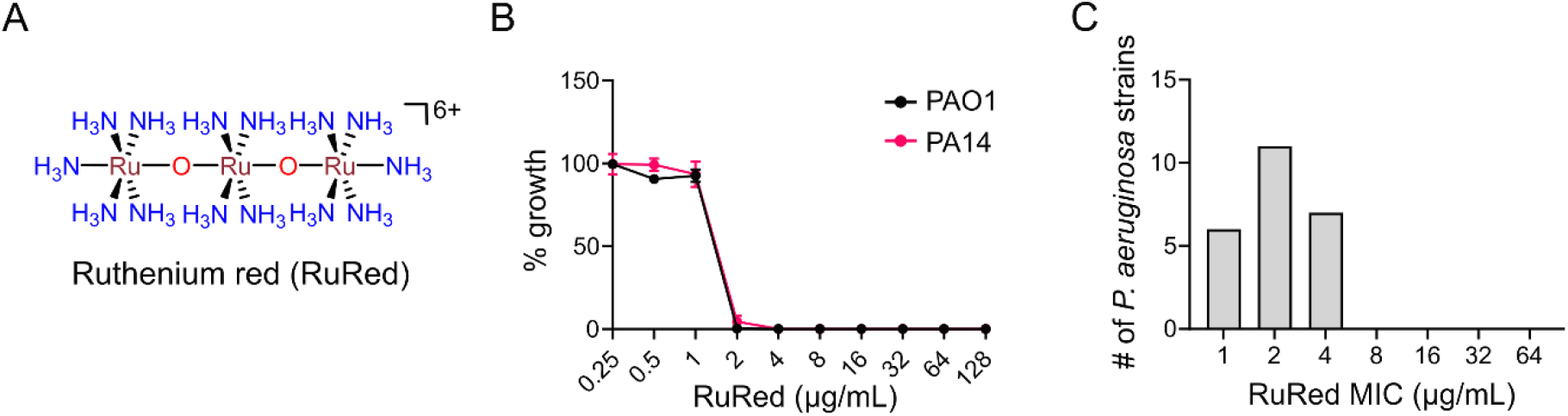
RuRed is antibacterial against *P. aeruginosa*. (A) Structure of Ruthenium red. (B) MIC curves for RuRed against laboratory strains of *P. aeruginosa* PAO1 and PA14 in MHB. (C) Distribution of RuRed MICs for a panel of *P. aeruginosa* isolates.

Given the potent activity of RuRed against *P. aeruginosa* PAO1 and PA14, we next tested a collection of 22 clinical isolates of *P. aeruginosa*. All isolates showed similar susceptibility to RuRed in MHB, with a MIC range of 1-4 µg/mL and a median of 2 µg/mL (Fig. 1C and Table S1). Testing of several other Gram-negative and –positive pathogens showed that, while most species had some level of RuRed susceptibility in MHB, *P. aeruginosa* strains were uniformly and uniquely highly susceptible to this compound (Table S1). We note that, while RuRed has been used experimentally for decades, its purity can vary between suppliers and is often not reported (25). Therefore, the purity of the commercially available RuRed used in this work was assessed by ultraviolet-visible (UV-vis) spectroscopy (Fig. S2). Our samples showed strong absorbance at 533 nm, characteristic of RuRed, and a second smaller peak at 260 nm, whose identity is not known but is often observed in commercial RuRed samples as a minor impurity (25). We also observed a minor peak at 360 nm which likely corresponds to the Ru360, a known impurity of RuRed preparations (26–28).

### Mutations in PA4394 (MscS-1) or PA4380 (ColS) decrease *P. aeruginosa* susceptibility to RuRed

To further characterize the antibacterial activity of RuRed against *P. aeruginosa*, we isolated spontaneous resistant mutants by plating the wild-type strain on an inhibitory concentration of RuRed. Mutants recovered using this method had a 4-fold increased MIC compared to the parental strain (Fig. 2A). Next, we sequenced the genomes of four mutants and the parental strain to identify any genetic differences. One of the four suppressor mutants (“suppressor1”) contained a mutation in the gene PA4394, which encodes a predicted mechanosensitive ion channel involved in osmotic tolerance called MscS-1 (29). This mutation, a point mutant at base 219, would result in a lysine to asparagine amino acid change (Fig. 2B). The other three mutants (collectively “suppressor2”) had an identical point mutation in the PA4380 gene at position 601, causing in a single amino acid change (Fig. 2B). PA4380 encodes the sensor kinase ColS that, together with the response regulator ColR encoded by PA4381, is part of the ColRS two-component sensor system (30, 31).

**Fig 2.**
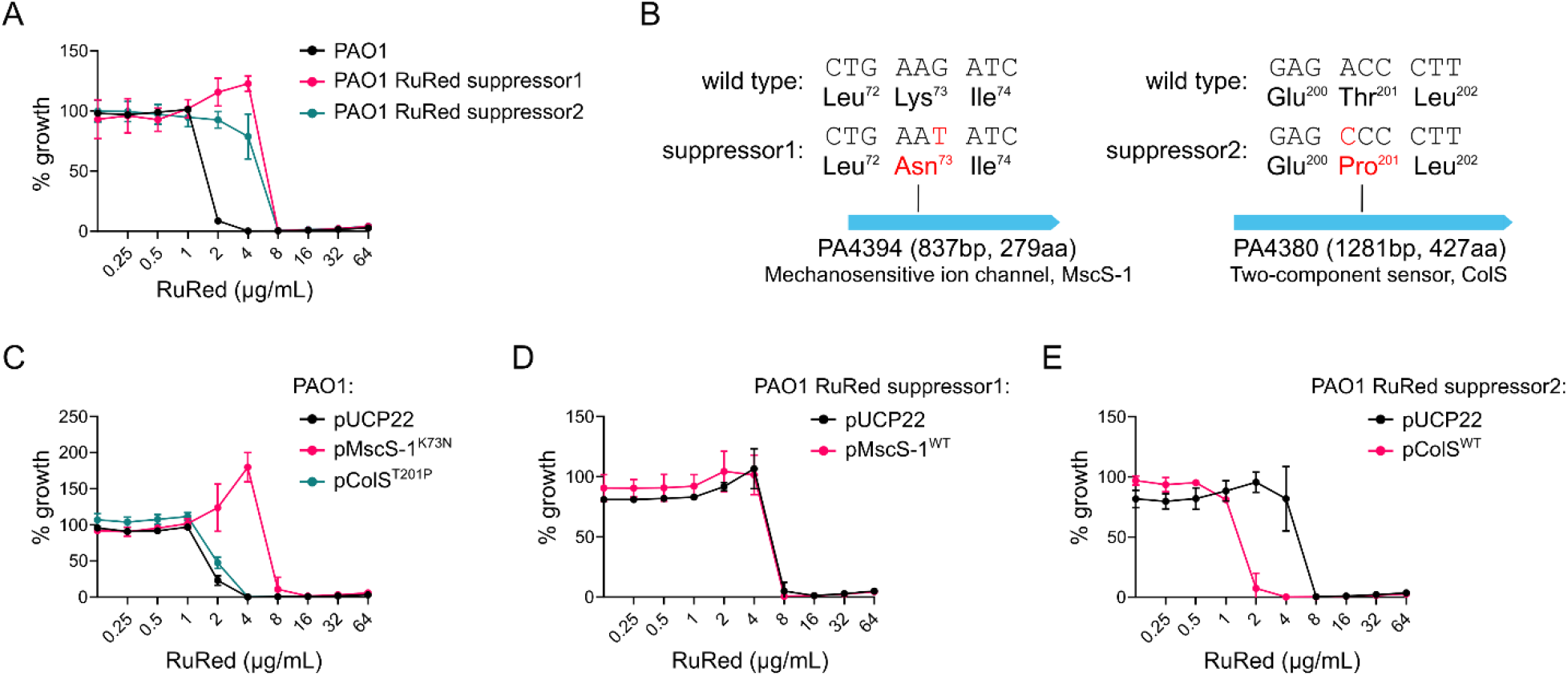
Genetic analysis of RuRed suppressors. (A) MICs of wild-type *P. aeruginosa* PAO1 or 2 independent suppressors. (B) Whole-genome sequencing of suppressors identified point mutations in *mscS-1* (suppressor1) or *colS* (suppressor2). (C) RuRed MICs of wild-type PAO1 containing vector control pUCP22 or expressing the mutant alleles of MscS-1 (pMscS-1^K73N^) or ColS (pColS^T201P^). (D) RuRed MICs of suppressor1 containing vector control pUCP22 or expressing the wild-type allele of MscS-1 (pMscS-1^WT^). (E) RuRed MICs of suppressor2 containing vector control pUCP22 or expressing the wild-type allele of ColS (pColS^WT^).

To confirm the role of MscS-1 and ColS in *P. aeruginosa* susceptibility to RuRed, the wild-type (MscS-1^WT^ and ColS^WT^) and mutant (MscS-1^K73N^ and ColS^T201P^) alleles were cloned into pUCP22 plasmid and expressed in the wild-type and mutant strains. Expression of MscS-1^K73N^, but not ColS^T201P^, in the wild-type background increased resistance of *P. aeruginosa* to RuRed (Fig. 2C). Interestingly, overexpression of the wild-type MscS-1 allele in the suppressor1 background did not restore susceptibility to RuRed (Fig. 2D), whereas overexpression of ColS^WT^ was able to restore sensitivity in the suppressor2 background (Fig. 2E). All constructs were sequence confirmed and produced similar levels of recombinant protein (Fig. S3). Thus, these results were not due to a lack of protein expression but instead suggest a more complex interaction between the wild-type and mutant alleles for each gene.

To further investigate the interactions of these alleles in RuRed susceptibility, we obtained insertionally-inactivated mutants in *mscS-1* and *colS* from the *P. aeruginosa* MPAO1 transposon mutant collection (32). We reasoned that this would allow us to determine the contribution of the wild-type or mutant alleles in a background devoid of a functional chromosomal copy of the genes. Unlike the suppressor mutants we isolated, both *mscS-1* and *colS* transposon mutants had identical RuRed MICs to the parental MPAO1 strain (Figure 3A). This indicates that the RuRed resistance seen in our suppressor mutants was not simply due to production of non-functional variants of these gene products. Expression of MscS-1^K73N^ increased RuRed resistance of both the MPAO1 wild-type (Figure 3B) and *mscS-1* transposon mutant background (Figure 3C), whereas expression of MscS-1^WT^ or the vector control had no effect. ColS^T201P^ expression had no effect in the wild-type background (Figure 3D), but significantly increased resistance in the *colS* mutant (Figure 3E) while ColS^WT^ had no impact in either background. Taken together, these results suggest that MscS-1^K73N^ is dominant to MscS-1^WT^, and ColS^WT^ is dominant to ColS^T201P^ for the observed RuRed resistance phenotypes. Given the fact that *mscS-1* and *colS* are not essential for *P. aeruginosa* viability, and the identified suppressor mutations impart a modest 4-fold increase in resistance to RuRed, it is unlikely that these gene products are the ultimate target of RuRed, but instead play an accessory role in susceptibility.

**Fig 3.**
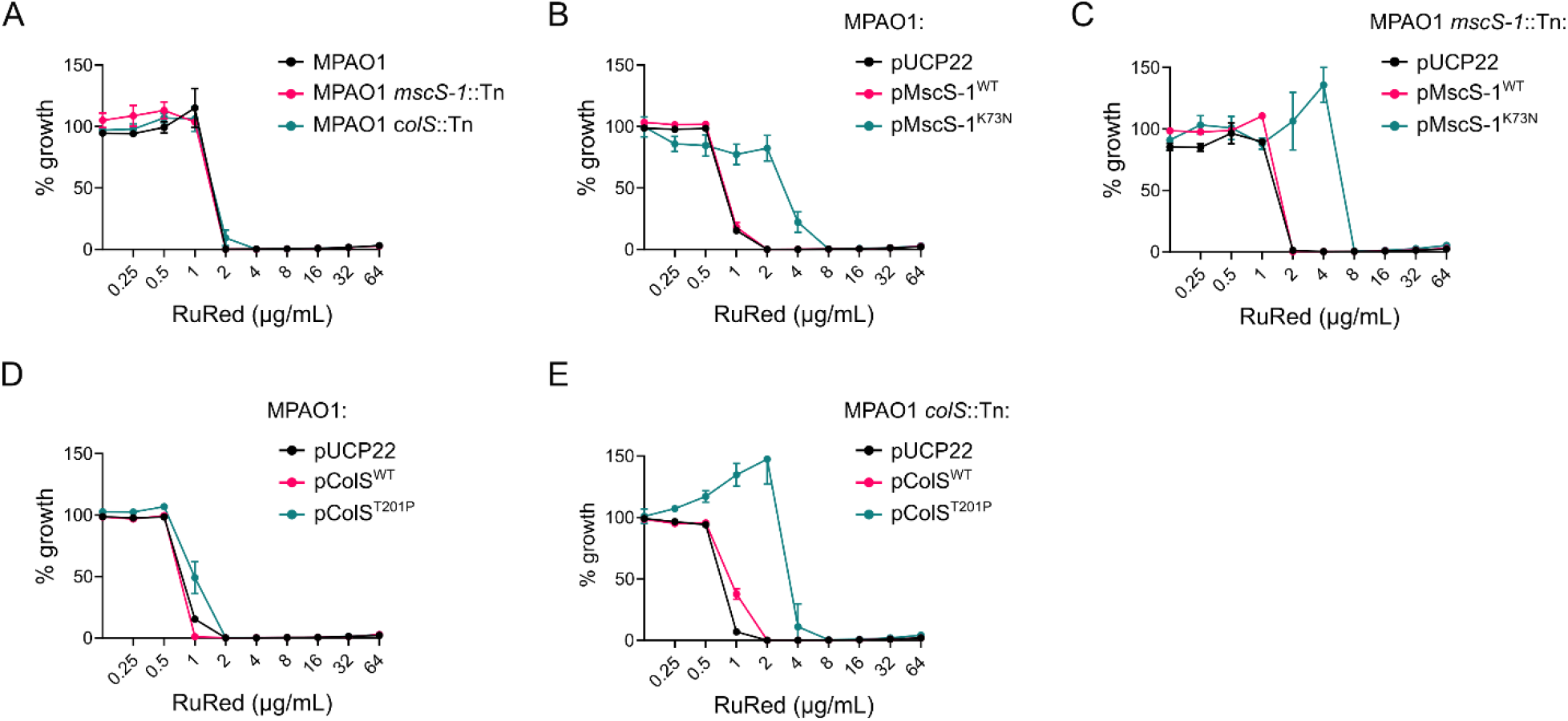
Effect of expression of MscS-1 and ColS alleles in wild-type and mutant backgrounds on RuRed MIC. (A) RuRed MIC curves of wild-type *P. aeruginosa* MPAO1 and MscS-1 and ColS transposon mutants. (B) RuRed MIC curves of wild-type MPAO1 containing vector control pUCP22 or expressing the WT or mutant allele of MscS-1. (C) RuRed MIC curves of MscS-1 transposon mutant containing vector control pUCP22 or expressing the WT or mutant allele of MscS-1. (D) RuRed MIC curves of wild-type MPAO1 containing vector control pUCP22 or expressing the WT or mutant allele of ColS. (E) RuRed MIC curves of ColS transposon mutant containing vector control pUCP22 or expressing the WT or mutant allele of ColS.

### RuRed analogs are antibacterial

In addition to RuRed, several related ruthenium compounds have been investigated for their ability to inhibit the MCU in eukaryotic cells (33, 34). Given their structural and functional similarity to RuRed, and our ability to synthesize pure compounds in-house, we tested the antibacterial activity of the well-characterized dinuclear ruthenium compounds Ru265, Ru270 and Ru360’ (Fig. 4A) (33). All 3 possessed antibacterial activity against *P. aeruginosa* PAO1, with MICs of 8, 64, and 4 µg/mL, respectively (Fig. 4B). In addition, all *P. aeruginosa* clinical isolates tested were susceptible to these compounds, with very similar MICs (Table S1). We also tested 6 mononuclear and 2 additional dinuclear Ru compounds, which had varying levels of antibacterial activity against *P. aeruginosa* PAO1 (Fig. S4). Like RuRed, Ru265, Ru270, and Ru360’ were less effective against non-*P. aeruginosa* bacterial species (Table S1). One exception, however, was the activity of Ru360’ against *Acinetobacter baumannii*. All tested *A. baumannii*, including common laboratory strains and clinical isolates, were susceptible to Ru360’ at concentrations similar to *P. aeruginosa* (Table S1).

**Fig 4.**
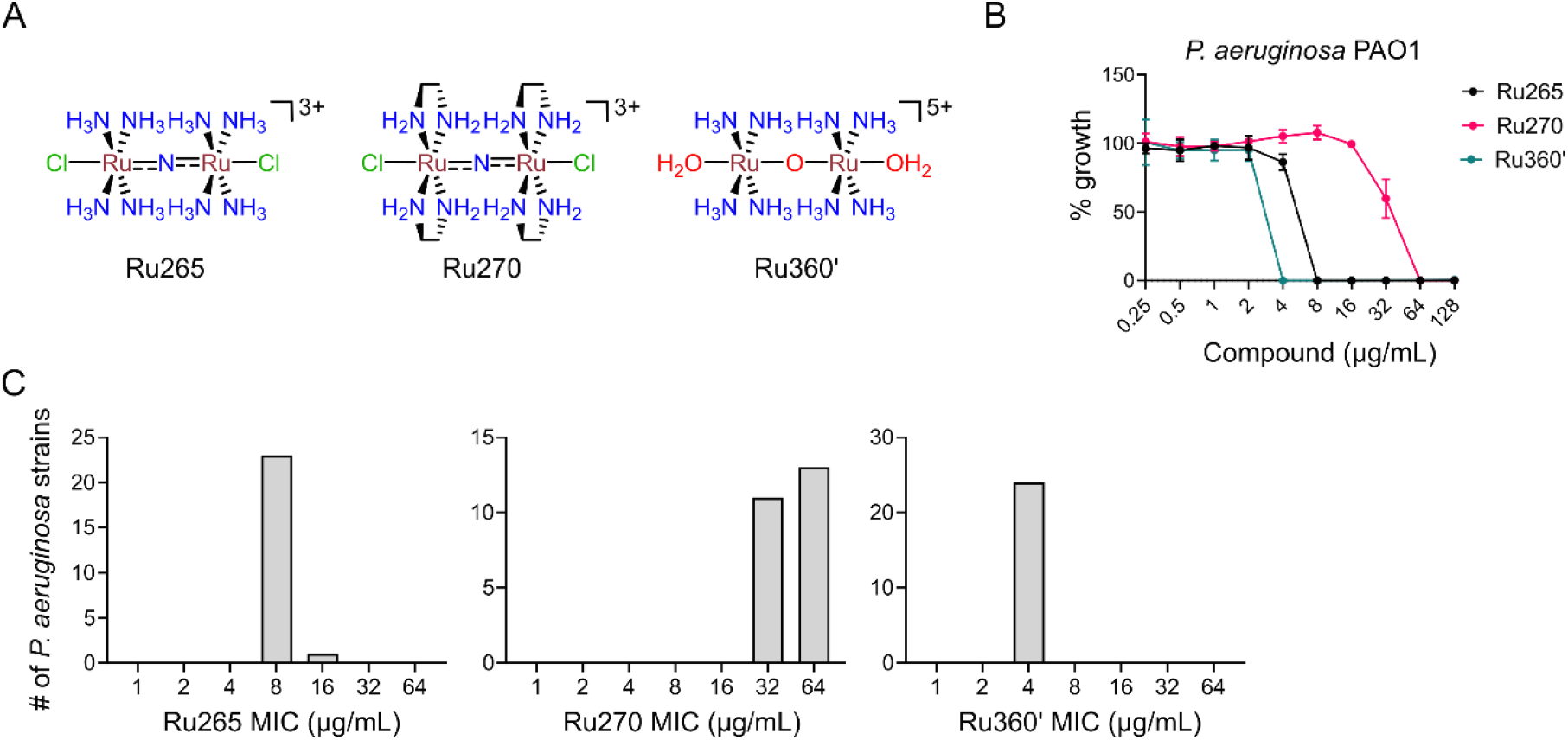
RuRed analogs Ru265, Ru270, and Ru360’ have antibacterial activity. (A) Structures of Ru255, Ru270 and Ru360’. (B) MIC curves of the indicated compounds against *P. aeruginosa* PAO1. (C) Distribution of MICs for a panel of *P. aeruginosa* isolates for the 3 compounds.

### Ruthenium compounds are effective in a *Galleria mellonella* infection model

Given the *in vitro* efficacy of the ruthenium compounds, in particular that of RuRed, Ru265, and Ru360’, we next used a *Galleria mellonella* infection model to assess their ability to treat infection *in vivo. G. mellonella* larvae are a common model system used to assess antibacterial activity, and activity in this model often translates to rodent models of infection (35). Groups of 10 larvae were infected with *P. aeruginosa* PAO1 and treated with a single dose of compound 15 min post-infection, then monitored for survival every 24h for 72h. Treatment of infected larvae with RuRed significantly enhanced survival of *G. mellonella* larvae in a dose-dependent manner, with 80% survival at the highest dose tested (Fig. 5A). Control larvae infected and treated with vehicle solution succumbed to infection by 24h, whereas those treated with the antibiotic ciprofloxacin survived for the duration of the experiment. Ru265 was similarly effective, with 90% survival at the highest dose tested (Fig. 5B). Treatment with Ru360’ also significantly extended survival compared to untreated larvae (Fig. 5C). However, treatment of uninfected larvae with Ru360’ revealed that this compound had toxic effects on its own, with the highest dose of Ru360’ resulting in the death of 70% of larvae (Fig. 5D). In contrast, the highest doses of RuRed and Ru265 had no negative effects on survival of uninfected larvae (Fig. 5D). Taken together, these results show that the *in vitro* efficacy of these compounds translates to an invertebrate model of infection.

**Fig 5.**
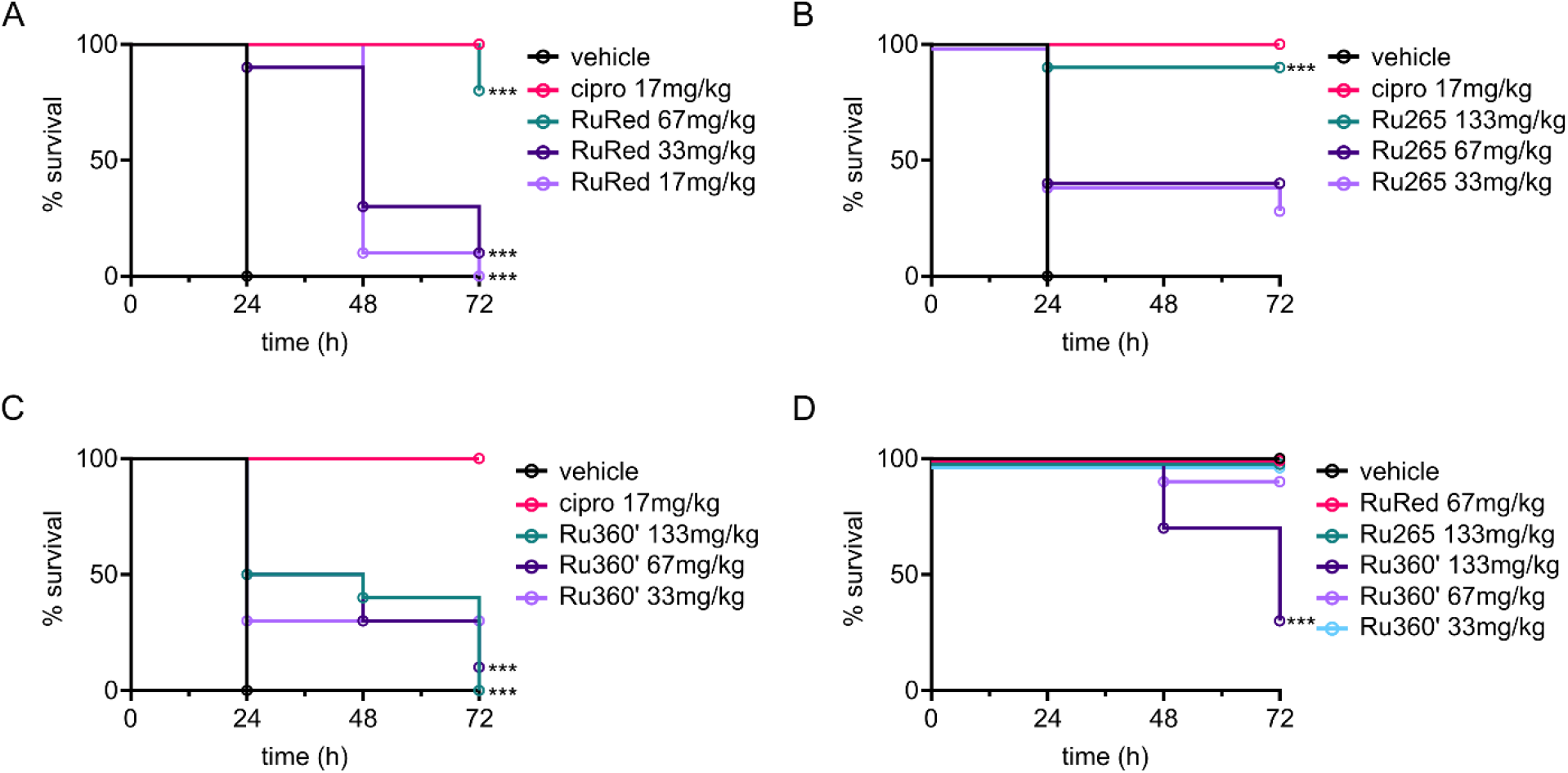
Ruthenium compounds are effective in a *G. mellonella* model of infection. Kaplan-Meier survival curves from *G. mellonella* larvae infected with *P. aeruginosa* PAO1 and treated with various doses of (A) RuRed, (B) Ru265, (C) Ru360’, or ciprofloxacin (cipro) or vehicle controls. (D) Uninfected larvae were injected with PBS and subsequently injected with the indicated compounds. In all experiments, survival was monitored every 24h for a total of 72h. *** indicates significant difference from the vehicle control. Significance was determined by using the Gehan-Breslow-Wilcoxon test with a Bonferroni correction for multiple comparisons.

### RuRed is effective in a *P. aeruginosa* mouse infection model

Our results with *G. mellonella* prompted us to evaluate RuRed in a mammalian infection model. We used a systemic mouse infection model, where mice were infected intraperitoneally with 10^6^ CFU/mL *P. aeruginosa* PAO1 and treated 1h and 5h post-infection with either 2mg/kg RuRed or vehicle control. Additionally, 4 infected mice were euthanized 1h-post infection to establish bacterial burden (colony forming units, CFUs) in the spleen, kidney, liver and bloodstream at the time of treatment (Fig. 6). Vehicle-treated mice had 1000 to 10000-fold increased bacterial CFU burden in the various organs and blood compared to the burden at the start of treatment (Fig. 6). In contrast, RuRed treated animals had either the same or slightly reduced bacterial CFUs compared to the initial burden seen 1h post-infection. Thus, RuRed shows potent antibacterial activity in a mouse model of systemic *P. aeruginosa* infection.

**Fig 6.**
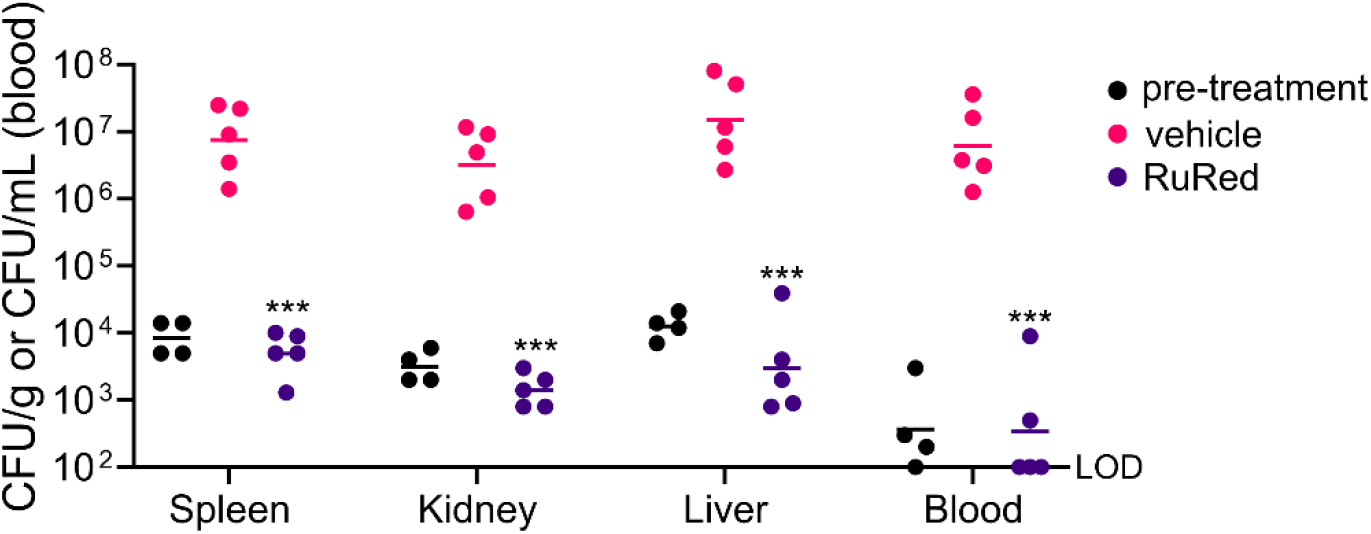
RuRed is effective in a systemic mouse infection model. Mice infected intraperitoneally with *P. aeruginosa* PAO1 were euthanized 1h post-infection for CFU determination (pre-treatment) or treated 1h and 5h post-infection with vehicle or RuRed (2mg/kg). Treated mice were euthanized 10h post-infection for CFU determination. Each point represents an individual animal, with the median indicated by the line. *** P<0.05 compared to vehicle control. Significance was determined using the Mann-Whitney test.

### Ruthenium compounds synergize with antibiotics *in vitro*

Although our *in vivo* studies show that these ruthenium complexes show promise as treatments for *P. aeruginosa* infection, it is also known that some of these compounds can have toxic side effects (36, 37). A common approach to ameliorate toxic side effects is to identify a partner compound that reduces the dose of the toxic molecule required for efficacy, resulting in a dose-sparing effect (38, 39). We tested combinations of RuRed with antibiotics in checkerboard assays and found synergy with several antibiotics normally used against Gram-positive bacteria, including rifampicin, vancomycin, novobiocin and erythromycin (Fig. S5 A-D). Similarly, Ru265, Ru270, and Ru360’ also synergized with rifampicin (Fig. S5 E and F), with Ru265 showing particularly strong synergy as evidenced by a fractional inhibitory concentration index (FICI) of 0.04 (Fig. 7A). Interestingly, this synergy also extended to *E. coli* (Fig. S6A). One possible explanation for these synergies is that the ruthenium compounds physically disrupt the outer membrane to allow entry of these antibiotics (40). To investigate this, we used a lysozyme lysis assay where physical disruption of the *E. coli* outer membrane allows lysozyme to reach the peptidoglycan layer and lyse the bacterial cells. Treatment with the control compound SPR741 led to nearly 60% lysis, whereas RuRed treatment resulted in no change in lysis compared to the no drug control (Fig. S6B). Thus, although RuRed can potentiate large scaffold antibiotics, the mechanism of synergy does not seem to be through outer membrane disruption.

**Fig 7.**
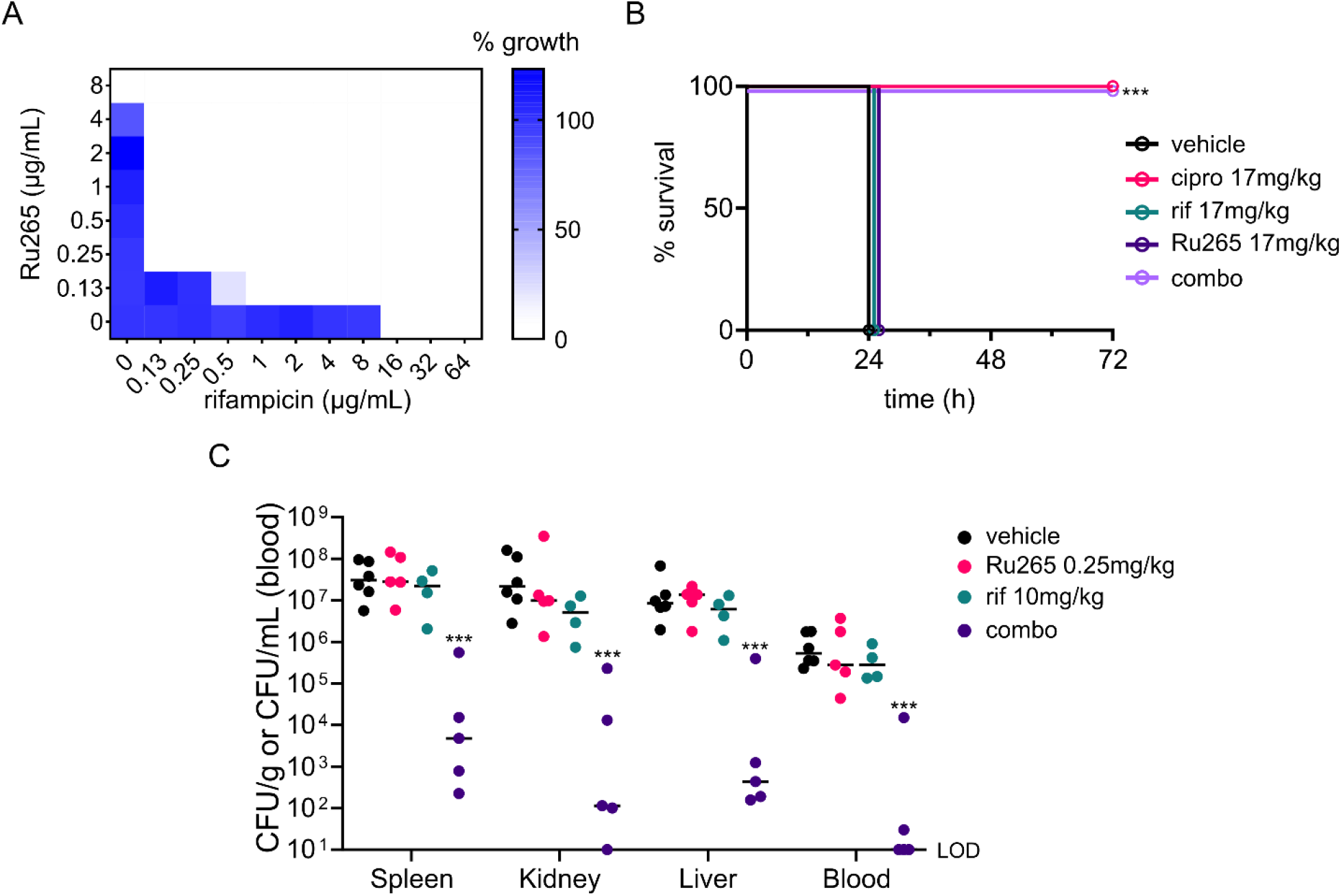
The combination of Ru265 and rifampicin is effective in vitro and in vivo. (A) Checkerboard assay showing synergistic effects (FICI=0.04) of Ru265 and rifampicin against *P. aeruginosa*. (B) *G. mellonella* larvae were infected with *P. aeruginosa* PAO1 and treated with monotherapies of rifampicin (rif) or Ru265, or the combination of both (combo). Control larvae were treated with ciprofloxacin (cipro) or vehicle. *** indicates significant difference from the vehicle control. Significance was determined by using the Gehan-Breslow-Wilcoxon test with a Bonferroni correction for multiple comparisons. (C) Systemic mouse infection model. Mice infected intraperitoneally with *P. aeruginosa* PAO1 were treated at 1h and 5h post-infection and were euthanized 10h post-infection for CFU determination. *** P <0.05 compared to vehicle control determine by Kruskal-Wallis test.

### The dose-sparing combination of RuRed or Ru265 and rifampicin is effective in vivo

The synergies of RuRed and Ru265 with rifampicin, and their favorable efficacy *in vivo*, encouraged us to test these combinations in the *G. mellonella* infection model. Larvae were infected with a lethal dose of *P. aeruginosa* and treated with the monotherapies or combinations. Treatment with sub-therapeutic levels of RuRed or rifampicin failed to cure infected larvae (Fig. S7). However, the treatment with the combination of RuRed and rifampicin resulted in 100% survival of the infected larvae, consistent with the observed *in vitro* synergies. Similarly, the combination of Ru265 and rifampicin completely cured infected larvae at doses where neither drug was effective on their own (Fig. 7B).

Next, we tested the combination of Ru265 and rifampicin in the systemic mouse infection model. We chose Ru265 due to the strong *in vitro* synergy with rifampicin seen in the checkerboard assays and efficacy in *G. mellonella*. Mice infected with *P. aeruginosa* and treated with the monotherapies had similar bacterial burdens in their blood and organs as vehicle-treated controls (Fig. 7C). Conversely, mice given the combination of Ru265 and rifampicin had a significant reduction in bacterial load in all organs and blood. These results show that addition of rifampicin can have a significant dose-sparing effect for Ru265 treatment, suggesting a promising approach to treat *P. aeruginosa* infection.

## Discussion

As antibiotic resistance continues to impact patients globally (41), innovative methods and approaches to drug discovery will be required to maintain the antibacterial pipeline (9). Recent studies have emphasized that metal complexes, which are often overlooked as potential antibiotics, can be productive sources of antibacterial and antifungal chemistry. Frei and colleagues screened nearly 1000 metal compounds for antimicrobial activity and found a hit-rate more than 10x higher than from conventional organic compound libraries, with many of these showing no apparent toxicity against mammalian cells (16). Gallium has shown promise as an antibacterial therapy, including in preliminary human trials (42, 43). A wide variety of ruthenium complexes have been investigated for antimicrobial activity, although there is limited *in vivo* efficacy data, particularly for Gram-negative bacteria (44–47). Our work shows that metal compounds like RuRed and Ru265 can be effective both *in vitro* and *in vivo* against *P. aeruginosa*, a notoriously difficult-to-treat Gram-negative bacterial pathogen.

*P. aeruginosa* showed a unique pattern of susceptibility to the polynuclear ruthenium compounds tested in this work. The reason(s) for the enhanced susceptibility of *P. aeruginosa* to the ruthenium compounds described here, compared to other bacteria, remain to be determined. In our previous work, we found that RuRed had no activity against *K. pneumoniae* in LB growth media but did have potent activity against *K. pneumoniae* grown in human serum (17). Curiously, MHB media seems to enhance the activity of RuRed, as we observed an MIC in this media even for *K. pneumoniae*. This suggests that growth media formulation influences susceptibility to these compounds. For example, serum is known to increase membrane permeability which enhances penetration of large scaffold drugs normally excluded by the outer membrane (48). We previously identified several *K. pneumoniae* gene knockouts that were susceptible to RuRed in LB media, such as those involved in outer membrane biogenesis and DNA repair pathways, and similar pathways have been shown to affect RuRed sensitivity in *E. coli*, further supporting the condition-specific antibacterial efficacy of RuRed (17, 49). It is possible that RuRed uses an entry mechanism into the cell that is constitutively active in *P. aeruginosa*, but only induced in other species under certain conditions. In any case, all bacterial species we tested were more resistant to the ruthenium compounds than *P. aeruginosa*, which indicates that there is some intrinsic property of *P. aeruginosa* that sensitizes them to these compounds. The one exception was our finding that Ru360’ had strong antibacterial activity against *A. baumannii* strains while RuRed, Ru265, and Ru270 had little or none. Overall, this suggests that this family of ruthenium compounds may have several possible modes of uptake into the cell, different mechanisms of action, or both. From a structural perspective, our data suggests that positive charge is important for activity of these ruthenium compounds, as the negatively charged dinuclear compounds tested were inactive (see Figure S3). However, a more thorough structure-activity-relationship campaign with a broader range of compounds should be a priority for future work.

We identified separate mutations in *mscS-1* and *colS* that led to a modest 4-fold increase in *P. aeruginosa* resistance to RuRed. However, elucidating the precise role that these loci play in resistance requires further work. It is likely that neither MscS-1 nor ColS are direct antibacterial targets of RuRed, as knockout mutants in both loci are viable (32). Experimental evidence has shown that MscS-1 is a mechanosensitive ion channelthat releases cytoplasmic osmolytes in response to osmotic stress (29). The MscS-1^K73N^ mutation we identified is unlikely to result in a loss of function; we showed that a *mscS-1* transposon mutant was just as susceptible as wild-type *P. aeruginosa*, and expression of MscS-1^K73N^ from a plasmid in the wild-type background led to resistance, suggesting the mutant allele is dominant. Interestingly, the eukaryotic target of RuRed and analogs are ion channels like the MCU, although it remains to be seen if RuRed directly interacts with MscS-1. The *colS* gene encodes the sensor histidine kinase ColS which, as part of the ColRS two-component system in various *Pseudomonas* species, has been implicated in virulence, host colonization, membrane integrity, and antibiotic resistance in certain genetic contexts (30, 50, 51, 31). The ColRS system regulates genes directly downstream from the *colRS* cluster, and these genes, in particular *warA* and *warB* have been shown to be involved in outer membrane modification and permeability (51–54). Therefore, we speculate that ColS^T201P^ results in dysregulation of the ColRS system, leading to changes in genes expression which in turn may prevent RuRed accumulation. Future studies to determine the mechanism of action of these ruthenium complexes should help uncover their ultimate targets and shed light on the unique susceptibility of *P. aeruginosa*.

We have shown that RuRed and Ru265 are effective *in vivo* therapies against *P. aeruginosa* infection in two different models. While the translation from *in vitro* to *in vivo* activity is clear, one of the major barriers to further developing these ruthenium compounds as antibiotics will likely be their toxicity. Indeed, we saw evidence of toxicity of Ru360’ in the *G. mellonella* infection model, and both RuRed and Ru265 have shown dose-dependent toxicity in mice (36, 37). Ideally, analogs could be developed that reduce toxicity; however, an alternative strategy is to use synergistic combinations of compounds. Combinations of antibiotics or small molecules that synergize can significantly reduce toxicity concerns due to the combinations’ dose-sparing effect (38, 55, 56). Indeed, we found that synergistic combinations of RuRed or Ru265 with rifampicin completely cured infected *G. mellonella*, and significantly reduced the bacterial burden in the mouse model. For the *G. mellonella* infections, the doses of the ruthenium compounds in the combination therapies were reduced nearly 10-fold compared to the monotherapy trials, and it is likely that with additional optimization this could be reduced further. The mechanism for the synergy seen with rifampicin requires further investigation, though our data suggest that it is not by physical disruption of the membrane. It is interesting to note that RuRed has been shown to bind to nucleic acids, which may suggest a possible route to synergy with an antibiotic like rifampicin (57). While we have not evaluated the toxicity of Ru265, we note that cell-based assays suggest minimal cytotoxicity of this compound (33), and thus Ru265 may be a promising lead compound for future development.

Much of the work on RuRed and analogs like Ru265, Ru270, and Ru360’ has focused on their ability to inhibit eukaryotic ion channels like the MCU (19). For antibacterial drug discovery, a compound with a eukaryotic target is often a liability. However, it was recently shown that inhibition of the MCU actually improves outcomes in CF models of *P. aeruginosa* infection (58). In that work, the authors showed that *P. aeruginosa* infection caused inflammation and impaired autophagy, which could be remedied by inhibition of the MCU with a compound called KB-R7943. In their CF mouse model, mice infected with *P. aeruginosa* and treated with this compound had significantly decreased bacterial burdens and increased survival. KB-R7493 is an organic compound that is structurally unrelated to the ruthenium compounds used here, and the authors’ ruled out any direct antibacterial activity caused by the KB-R7493 itself. Thus, it is tempting to speculate that in a CF model of *P. aeruginosa* infection, administration of compounds like RuRed or Ru265 could provide a dual benefit: 1) inhibition of the MCU, resulting in physiological corrections that improve host clearance of *P. aeruginosa* and 2) direct antibacterial activity against *P. aeruginosa*.

Overall, we have uncovered a series of compounds that are effective therapies against one of the most troublesome Gram-negative pathogens. Our work provides a framework for future studies that unravel the antibacterial mechanism and full *in vivo* potential of these promising ruthenium complexes.

## Materials and Methods

### Bacterial strains, media, and compounds

The bacterial strains used in this work are listed in Table S2. Strains were routinely cultured and maintained in LB (per liter: 10 g trypticase peptone, 5 g yeast extract, 10 g NaCl) or LB agar at 37°C. Cation-adjusted Mueller-Hinton II broth (MHB, BD, Cat. # 212322) was used for susceptibility testing. RuRed was purchased from Sigma-Aldrich (Cat. # R2751), and Ru265, Ru270 and Ru360’, and other ruthenium complexes were synthesized as previously described (33, 34, 59–66). All compounds were dissolved in water.

### Ultraviolet-visible (UV-vis) Spectroscopy of RuRed

Three samples of RuRed, from two different lot numbers but all separately ordered from the vendor, were analyzed via UV-vis spectroscopy to assess purity. Each RuRed sample (2.0 mg; 0.0024 mmol) was added to 10 mL 0.1 M ammonium acetate (NH_4_OAc) solution to give a 250 µM concentration, assuming the material was 100% pure. The solutions were then further diluted 10-fold in 0.1 M NH_4_OAc to an assumed final concentration of 25 µM. UV-vis spectra were taken at 25 °C using a Shimadzu UV-1900 Spectrophotometer (Shimadzu, Kyoto, Japan) fitted with a temperature-controlled circulating water bath. Purity of each sample was assessed by the magnitude of absorbance at 533 nm, characteristic of RuRed (25).

### Antibacterial MIC testing and checkerboard assays

Susceptibility MIC testing and checkerboards were performed using a broth microdilution assay in MHB, unless otherwise noted. Strains were plated on agar and a loopful of colonies was resuspended in 0.85% saline and adjusted to an OD_600nm_ of 0.5, or ∼7.5×10^8^ CFU/mL. For MIC testing, 0.67 µL of this suspension was added per mL of MHB, giving a final concentration of ∼5×10^5^ CFU/mL, and added to 96-well plates containing the serially diluted compounds of interest to give a final volume of 100 µL. Baseline OD_600nm_ reads were taken and plates were incubated stationary for 18h at 37°C and then OD_600nm_ was again measured using a plate reader. For MIC dose-response curves or checkerboard figures, growth was determined by subtracting baseline reads, and normalizing the no compound, solvent control to 100%. Reported MICs were determined by visual inspection of plates and reporting the lowest concentration at which no growth was visible. For checkerboards, the FIC of each compound was determined to be its MIC in combination with the other compound divided by its MIC alone. If a compound did not have an MIC, the next highest two-fold dilution to that tested was used. The reported FIC indices (FICIs) are the sum of FICs for the two compounds being tested (67). FICI values ≤ 0.5 were considered synergistic. MICs in serum were done as above but *P. aeruginosa* PAO1 was inoculated into 50% human serum (BioIVT), diluted using 1x M9 salts with 25 mM bicarbonate and 1x final concentration Biolog Redox Dye Mix A (Biolog, Cat.# 74221). This was added to 96-well plates containing the serially diluted compounds of interest to give a final volume of 200 µL, similar to previous work (17). MICs reported are from biological triplicate data.

### Suppressor generation and sequencing

Two biological replicates of *P. aeruginosa* PAO1 were grown overnight in LB, a sample of each was saved for genome sequencing, and 100µL of culture was plated onto LB agar plates containing 125µg/mL of RuRed. Two colonies from each biological replicate that grew on the RuRed plates were picked and struck onto LB agar. A single colony from each was inoculated in liquid culture and grown overnight, then genome prepped and cryostocked, and also verified for increased MIC to RuRed in MHB. Genomic DNA from the wild-type PAO1 biological replicates and 2 mutants from each replicate (6 samples total) were genome sequenced at the Microbial Genome Sequencing Center (MiGS, Pittsburgh, PA) with Illumina NextSeq 2000 platform and variant called using BreSeq (68) to identify mutations.

### Cloning and expression of PA4394 (mscS-1) and PA4380 (colS) alleles

Genomic DNA from wild-type *P. aeruginosa* PAO1 was used to PCR amplify the *mscS-1*^*WT*^ and *colS*^*WT*^ genes, and genomic DNA from the RuRed suppressors was used to PCR amplify *mscS-1*^*K73N*^ *and colS*^*T201P*^. *mscS-1*^WT^ and *mscS-1*^K73N^ were amplified with the primer pair 5’ atatggatccatggaattgaactacgaccgactgg 3’ and 5’ ataaagctttcagtggtgatggtgatgatggtcggccatcgcgccctgc 3’, which incorporated BamHI and HindIII restriction sites and a C-terminal histidine tag, digested and ligated into pUCP22 digested with BamHI and HindIII. *colS*^WT^ and *colS*^T201P^ were amplified with the primer pair 5’ atatggatccatggagtataagcagagcctcg 3’ and 5’ atagtcgactcagtggtgatggtgatgatgagcaacatcgagtaaaacttcg 3’, which incorporated BamHI and SalI restriction sites and a C-terminal 6x histidine tag, digested and ligated into pUCP22 digested with BamHI and SalI. After ligation, the constructs were electroporated into *E. coli* Top10 cells and plated on LB agar with 30 µg/mL gentamicin. Transformants were picked from plates, grown in liquid culture, and plasmids prepped for sequencing and subsequent transformation into *P. aeruginosa* PAO1 strains. Protein expression in normalized whole cell samples of *P. aeruginosa* strains overexpressing the proteins was verified by Western blot. Briefly, after SDS-PAGE and transfer to a nitrocellulose membrane, recombinant proteins were detected using primary anti-His antibody (ThermoFisher, Cat. # PA1-983B) and an anti-RNAP antibody (BioLegend, Cat. # 663104) as a loading control, followed by the secondary antibodies IRDye 800CW Goat anti-Rabbit and IRDye 680RD Goat anti-Mouse (LiCor, Cat. #925-32211 and Cat. #925-68070, respectively).

### G. mellonella infections

*G. mellonella* were reared in-house and larvae weighing approximately 300mg were used for all experiments. 100 µl of an overnight culture of *P. aeruginosa* PAO1 was added into fresh media and grown to and OD_600_ of 0.8, then washed twice in PBS and resuspended to a final density of 1-1.5×10^4^ CFU/mL. 10 µl of this was injected into the last proleg of the larvae using a syringe (U-100 Insulin Syringes, Cat. # 329461, BD) loaded into a syringe pump (KDS-100, KD Scientific) (69), for a total dose of 100-150 CFU of *P. aeruginosa* PAO1 per larvae. Uninfected controls were injected with PBS. After 15min, larvae were given a second injection of 10 µl containing drug or vehicle (water) control and incubated at 37°C. Survival was monitored every 24h for a total of 72h, and larvae were recorded as dead if they failed to respond to touch. Statistics were performed using GraphPad prism. The statistical significance of ruthenium compound treatments compared to vehicle control was determined using the Gehan-Breslow-Wilcoxon test with a Bonferroni correction for multiple comparisons. Using a defined significance threshold of 0.05, the Bonferroni-corrected significance threshold was calculated by dividing 0.05 by the number of comparisons. In most cases, this was 3, yielding a significance threshold of 0.017.

### Mouse studies

All mouse studies were conducted according to guidelines set by the Canadian Council on Animal Care using protocols approved by the Animal Review Ethics Board at McMaster University under Animal Use Protocol #20-12-43. All animal studies were performed with 6-10-week-old female CD-1 mice (Envigo). Female mice were used following previously established models, as well as ease of housing and randomization. No animals were excluded from analyses, and no sample or effect size assumptions were made. Sample sizes were selected based on results from pilot experiments.

CD-1 mice were infected intraperitoneally with ∼1×10^6^ CFU/mL of P. aeruginosa PAO1 with 5% porcine mucin. For the monotherapy, treatment with RuRed (2 mg/kg) or a vehicle solution (water) was administered intraperitoneally at 1h and 5h post-infection. For the combination therapy experiment, Ru265 (0.25 mg/kg), rifampicin (10 mg/kg), vehicle (water), or the combination was administered intraperitoneally at 1h post-infection. Due to the unknown pharmacokinetics of Ru265, a second treatment of Ru265 (0.25 mg/kg) was administered to the Ru265 monotherapy and combination therapy animals at 5h post-infection, and the other groups (vehicle and rifampicin monotherapy) received vehicle (water) injections. Blood was collected in Li heparin tubes by facial bleed 10h post-infection. Mice were then euthanized and the spleen, kidneys, and liver were collected into PBS at necropsy. Organs were homogenized using a high-throughput tissue homogenizer, serially diluted in PBS, and plated onto solid LB. Plates were incubated overnight at 37°C, and colonies were quantified to determine organ load. Statistics were performed using GraphPad Prism software. Significance was determined by the Mann-Whitney tests and a significance threshold of 0.05 for the RuRed experiments, and the Kruskal-Wallis test for the combination therapy experiments with a significance threshold of 0.05.

### Lysozyme assay for outer membrane permeability

Assay was performed as previously described (40) with modifications. *E. coli* BW25113 was grown overnight (∼18h) in MHB and subcultured 1:50 in fresh LB media at 37 °C with shaking at 250 rpm to mid-log (OD_600_ ∼0.4-0.5). Subcultures were centrifuged and resuspended in 5 mM HEPES pH 7.2 to OD_600_ ∼0.5. A volume of 1 mL of cells was added to microfuge tubes containing the indicated concentrations of compound and 50 µg/mL lysozyme (Sigma-Aldrich). Tubes were gently inverted to mix and left at room temperature for ∼30 min for any cell debris to settle. A volume of 200 µL was transferred from each tube to a clear, flat bottom 96-well plate, and OD600 was read using a Tecan Infinite M1000 Pro plate reader. Controls were included to ensure that the drugs alone did not lyse cells and to find the baseline levels of cell lysis by lysozyme alone.

## Acknowledgements

We thank Dr. J. Dennis and Dr. J. McCutcheon (University of Alberta) for providing plasmid pUCP22 and *P. aeruginosa* PAO1 and PA14 strains, Tim Cho and members of the Brown and Raivio laboratories for thoughtful suggestions and discussion of this work, and Dr. Gerry Wright for providing clinical isolates. Work in the laboratory of E.D.B. was supported by a Tier I Canada Research Chair award, a Foundation grant from the Canadian Institutes of Health Research (CIHR; FRN 143215), and a grant from the Ontario Research Fund (RE09-047). Work in the laboratory of J.J. Wilson was supported by NSF CAREER Award No. CHE-1750295. Work in the laboratory or T.R. was supported by operating grants from The National Sciences and Engineering Research Council (NSERC RGPIN-2021-02710) and The Canadian Institutes of Health Research (CIHR MOP 142347), and a project grant from the AMR-One Health Consortium, funded by the Major Innovation Fund program of the Ministry of Jobs, Economy and Innovation, Government of Alberta. B.S.W. was supported by a CIHR Postdoctoral Fellowship. J.J. Woods was supported by the National Science Foundation (DGE-1650441) and the American heart Association (20PRE35120390). All research was performed prior to K.K. beginning employment at Health Canada (HC). HC was not involved in the research.

