## Supplemental Figures and Tables for "Polynuclear ruthenium complexes are effective antibiotics against *Pseudomonas aeruginosa*"

### Supplementary Figures and Tables

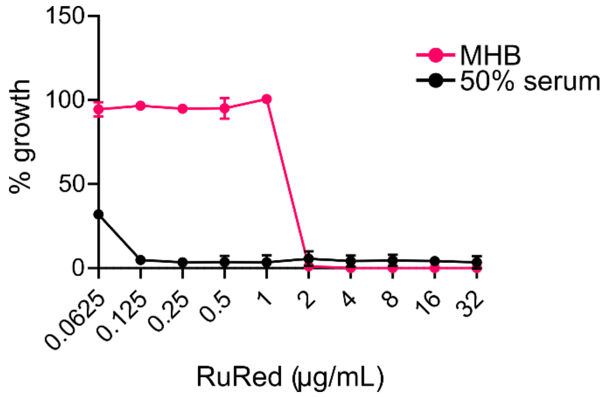

**Fig S1. Serum enhances activity of RuRed against *P. aeruginosa*.** MIC curves of RuRed against *P. aeruginosa* PAO1 grown in MHB and 50% human serum.

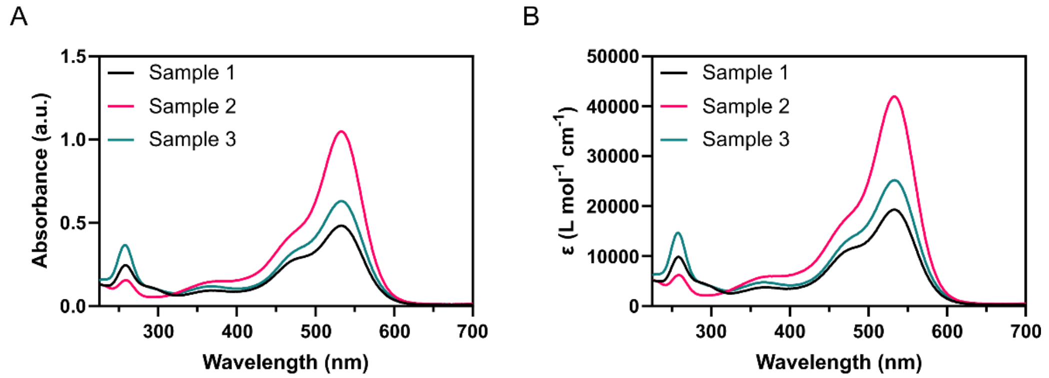

**Fig S2. Analysis of RuRed purity.** Three samples of RuRed (Sample 1-3) purchased from Sigma-Aldrich were analyzed by UV-vis spectroscopy. (A) Absorbance spectra for each sample and (B) molar extinction coefficients. Each sample was from a different vial of purchased RuRed; samples 1 and 3 were from the same lot number.

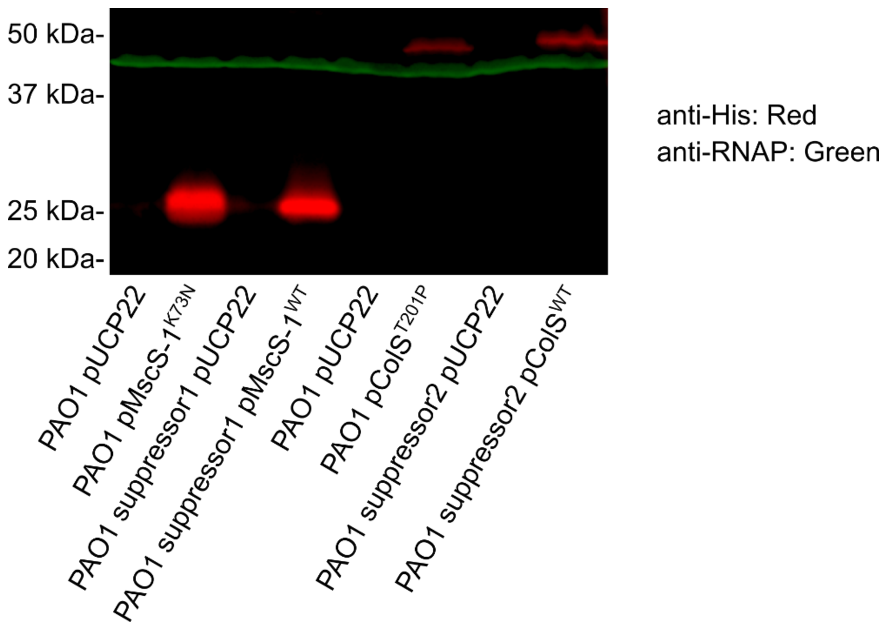

**Fig S3. Verification of recombinant protein expression by Western blot.** Histidine tagged $PA4394^{WT}$, $PA4394^{K73N}$, $PA4380^{WT}$, $PA4380^{T201P}$ were expressed in *P. aeruginosa* PAO1 from the pUCP22 plasmid and whole cell lysates were probed for expression of the recombinant proteins with an anti-His antibody (Red). The anti-RNAP antibody (green) was used as a loading control.

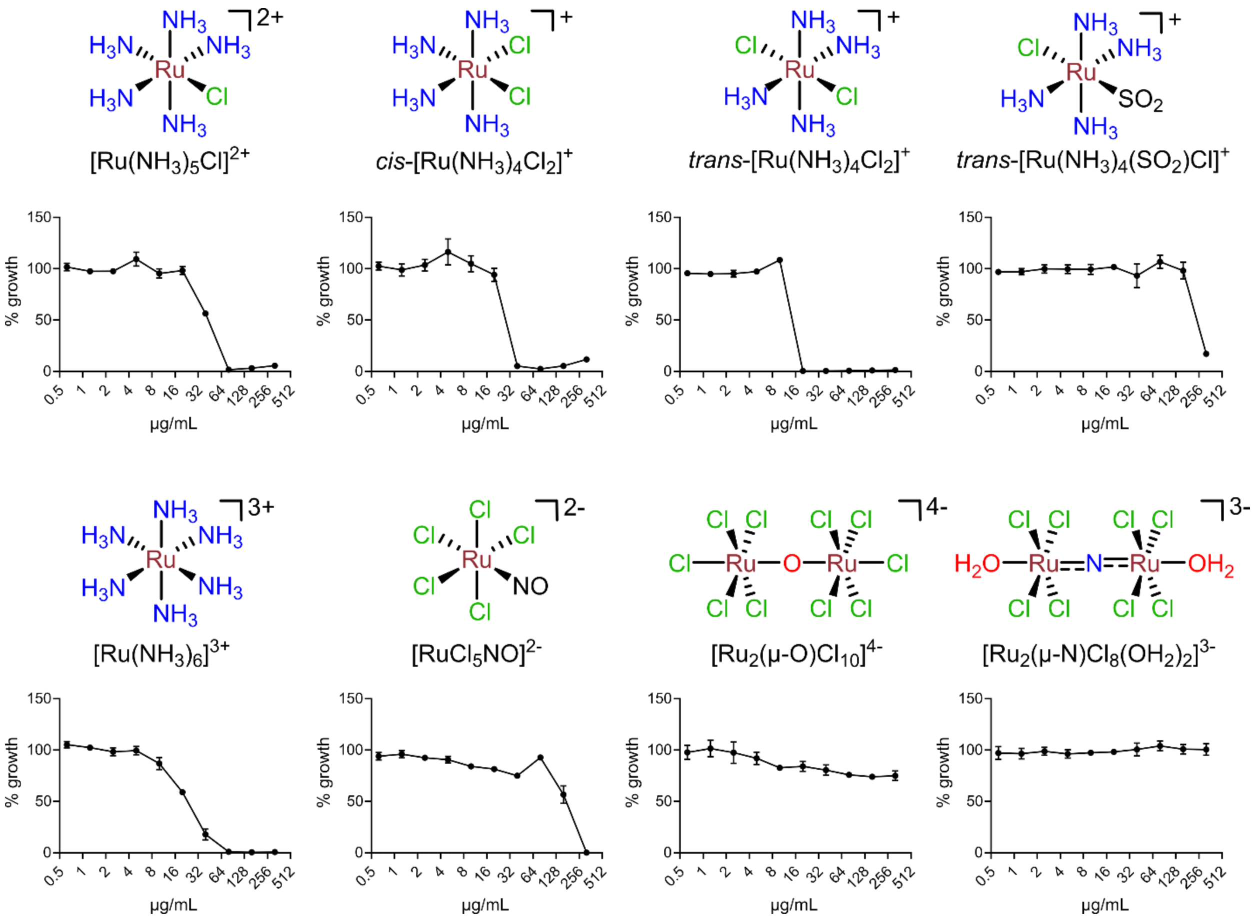

**Fig S4. Antibacterial activity of various mononuclear and dinuclear ruthenium compounds.** MIC curves for each compound against *P. aeruginosa* PAO1 in MHB.

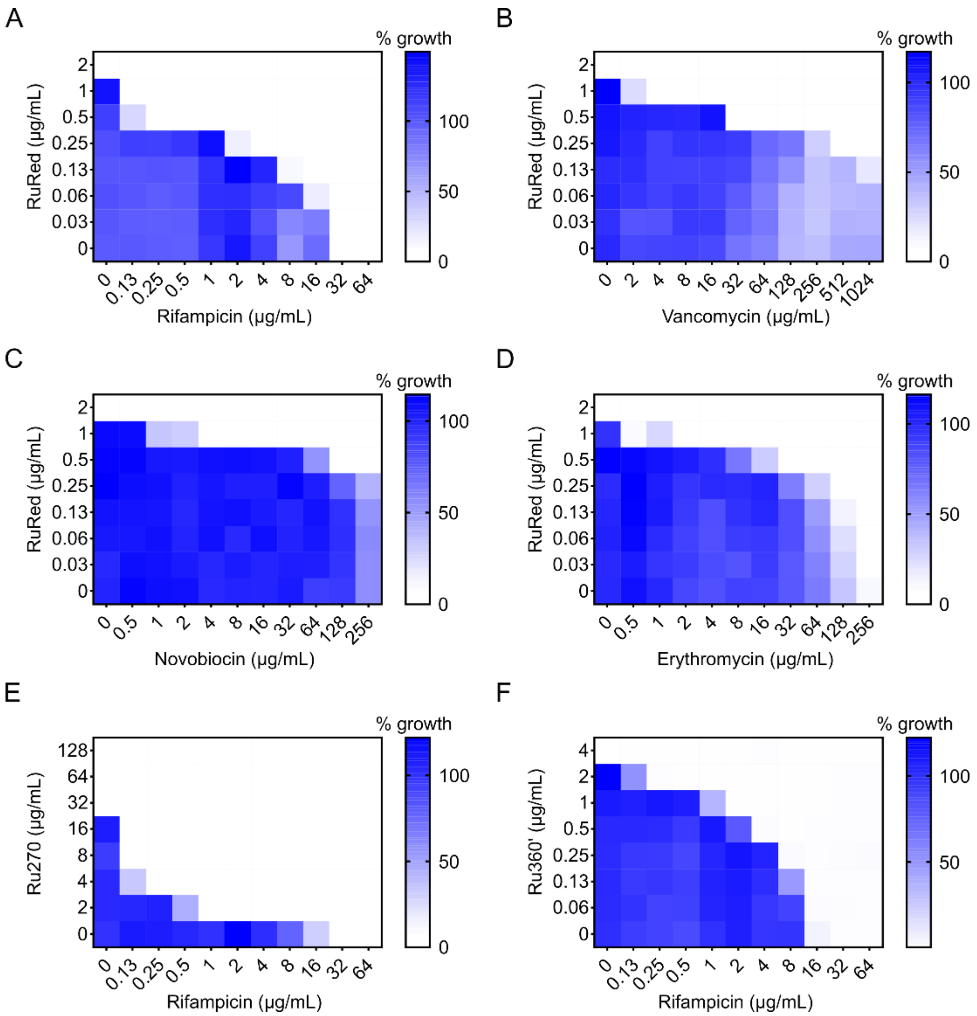

**Fig S5. Checkerboard assays to detect interactions between ruthenium compounds and antibiotics**. *P. aeruginosa* PAO1 susceptibility to RuRed paired with (A) rifampicin (FICI=0.25), (B) vancomycin (FICI=0.27), (C) novobiocin (FICI=0.5) and (D) erythromycin (FICI=0.31). All combinations were found to be synergistic against *P. aeruginosa* PAO1 (FICI≤0.5). (E) Ru270 paired with rifampicin (FICI=0.09) and (F) Ru360' paired with rifampicin (FICI=0.31). Both Ru270 and Ru360' show synergy with rifampicin.

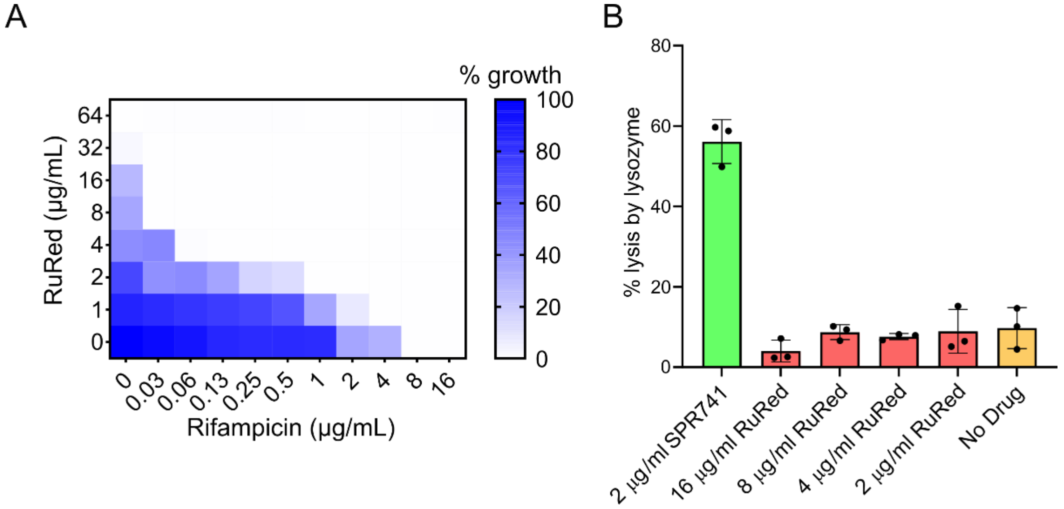

**Fig S6. RuRed does not cause outer membrane permeability.** (A) Checkerboard showing synergy between RuRed and rifampicin against *E. coli* BW25113 (FICI=0.13). (B) Lysozyme lysis assay. *E. coli* cells were treated with the indicated concentrations of each compound in the presence of lysozyme and assayed for lysis. The compound SPR741 served as a positive outer membrane permeabilizer and lysis control. RuRed showed no difference from the no drug controls.

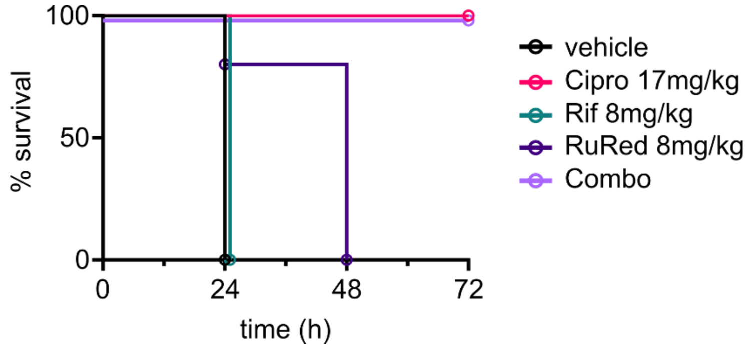

**Fig S7. The combination of RuRed and rifampicin is effective at treating *G. mellonella* larvae.** *G. mellonella* larvae were infected with *P. aeruginosa* PAO1 and treated with monotherapies of rifampicin (Rif) or RuRed, or the combination of both (combo). Control larvae were treated with ciprofloxacin (cipro) or vehicle.

**Table S1- MICs of ruthenium compounds**

| Strain | RuRed MIC* | Ru265 MIC | Ru360' MIC | Ru270 MIC |
|---|---|---|---|---|
| *Pseudomonas aeruginosa* PAO1 | 2 (8)* | 8 | 4 | 64 |
| *Pseudomonas aeruginosa* PA14 | 2 | 8 | 4 | 64 |
| *Pseudomonas aeruginosa* C0028 | 1 | 8 | 4 | 32 |
| *Pseudomonas aeruginosa* C0029 | 1 | 8 | 4 | 64 |
| *Pseudomonas aeruginosa* C0070 | 4 | 8 | 4 | 64 |
| *Pseudomonas aeruginosa* C0071 | 2 | 8 | 4 | 64 |
| *Pseudomonas aeruginosa* C0089 | 2 | 16 | 4 | 64 |
| *Pseudomonas aeruginosa* C0090 | 1 | 8 | 4 | 32 |
| *Pseudomonas aeruginosa* C0100 | 4 | 8 | 4 | 64 |
| *Pseudomonas aeruginosa* C0129 | 1 | 8 | 4 | 32 |
| *Pseudomonas aeruginosa* C0135 | 2 | 8 | 4 | 32 |
| *Pseudomonas aeruginosa* C0156 | 1 | 8 | 4 | 32 |
| *Pseudomonas aeruginosa* C0176 | 2 | 8 | 4 | 64 |
| *Pseudomonas aeruginosa* C0177 | 1 | 8 | 4 | 32 |
| *Pseudomonas aeruginosa* C0189 | 2 | 8 | 4 | 64 |
| *Pseudomonas aeruginosa* C0190 | 2 | 8 | 4 | 32 |
| *Pseudomonas aeruginosa* C0200 | 4 | 8 | 4 | 64 |
| *Pseudomonas aeruginosa* C0201 | 2 | 8 | 4 | 32 |
| *Pseudomonas aeruginosa* C0202 | 4 | 8 | 4 | 64 |
| *Pseudomonas aeruginosa* C0203 | 4 | 8 | 4 | 32 |
| *Pseudomonas aeruginosa* C0204 | 2 | 8 | 4 | 64 |
| *Pseudomonas aeruginosa* C0213 | 4 | 8 | 4 | 64 |
| *Pseudomonas aeruginosa* C0214 | 4 | 8 | 4 | 32 |
| *Pseudomonas aeruginosa* C0215 | 2 | 8 | 4 | 32 |
| *Acinetobacter baumannii* C0015 | 64 | 64 | 2 | >128 |
| *Acinetobacter baumannii* B80510 | 32 | 32 | 4 | >128 |
| *Acinetobacter baumannii* C0044 | 32 | 32 | 2 | >128 |
| *Acinetobacter baumannii* C0074 | >128 | 64 | 4 | >128 |
| *Acinetobacter baumannii* C0092 | >128 | 64 | 4 | >128 |
| *Acinetobacter baumannii* C0102 | 128 | 64 | 4 | >128 |
| *Acinetobacter baumannii* ATCC 19606 | 128 | 32 | 2 | 128 |
| *Escherichia coli* C0002 | 64 | 16 | 32 | >128 |
| *Serratia marcescens* C0016 | 64 | >64 | 128 | >128 |
| *Proteus mirabilis* C0027 | 64 | >64 | 32 | >128 |
| *Enterobacter cloacae* C0035 | 128 | 64 | 128 | >128 |
| *Klebsiella oxytoca* C0036 | 64 | 32 | 32 | >128 |
| *Klebsiella aerogenes* C0045 | 16 | 32 | 64 | >128 |
| *Morganella morganii* C0052 | 128 | >64 | 64 | >128 |
| *Citrobacter freundii* C0055 | 128 | 32 | 128 | >128 |
| *Serratia odorifera* C0082 | 64 | >64 | 32 | >128 |
| *Stenotrophomonas maltophilia* C0101 | 128 | >64 | 128 | >128 |
| *Citrobacter amalonaticus* C0120 | 64 | 16 | 64 | 128 |
| *Staphylococcus aureus* TCH1516 | 64 | 64 | >128 | >128 |
| *Klebsiella pnuemoniae* ATCC 43816 | 128 | 32 | 64 | >128 |
| *Klebsiella pneumoniae* MKP103 | 64 (>256)* | 64 | 16 | >128 |
| *Escherichia coli* BW25113 | 64 | 16 | 32 | 128 |

*- MICs in brackets are from experiments done in LB

### Table S2- Strains and plasmids used in this work

| Strain | Source/Notes |
|---|---|
| *Pseudomonas aeruginosa* PAO1 | Laboratory strain |
| *Pseudomonas aeruginosa* PA14 | Laboratory strain |
| *Pseudomonas aeruginosa* C0028 | McMaster clinical isolate |
| *Pseudomonas aeruginosa* C0029 | McMaster clinical isolate |
| *Pseudomonas aeruginosa* C0070 | McMaster clinical isolate |
| *Pseudomonas aeruginosa* C0071 | McMaster clinical isolate |
| *Pseudomonas aeruginosa* C0089 | McMaster clinical isolate |
| *Pseudomonas aeruginosa* C0090 | McMaster clinical isolate |
| *Pseudomonas aeruginosa* C0100 | McMaster clinical isolate |
| *Pseudomonas aeruginosa* C0129 | McMaster clinical isolate |
| *Pseudomonas aeruginosa* C0135 | McMaster clinical isolate |
| *Pseudomonas aeruginosa* C0156 | McMaster clinical isolate |
| *Pseudomonas aeruginosa* C0176 | McMaster clinical isolate |
| *Pseudomonas aeruginosa* C0177 | McMaster clinical isolate |
| *Pseudomonas aeruginosa* C0189 | McMaster clinical isolate |
| *Pseudomonas aeruginosa* C0190 | McMaster clinical isolate |
| *Pseudomonas aeruginosa* C0200 | McMaster clinical isolate |
| *Pseudomonas aeruginosa* C0201 | McMaster clinical isolate |
| *Pseudomonas aeruginosa* C0202 | McMaster clinical isolate |
| *Pseudomonas aeruginosa* C0203 | McMaster clinical isolate |
| *Pseudomonas aeruginosa* C0204 | McMaster clinical isolate |
| *Pseudomonas aeruginosa* C0213 | McMaster clinical isolate |
| *Pseudomonas aeruginosa* C0214 | McMaster clinical isolate |
| *Pseudomonas aeruginosa* C0215 | McMaster clinical isolate |
| *Acinetobacter baumannii* C0015 | McMaster clinical isolate |
| *Acinetobacter baumannii* B80510 | McMaster clinical isolate |
| *Acinetobacter baumannii* C0044 | McMaster clinical isolate |
| *Acinetobacter baumannii* C0074 | McMaster clinical isolate |
| *Acinetobacter baumannii* C0092 | McMaster clinical isolate |
| *Acinetobacter baumannii* C0102 | McMaster clinical isolate |
| *Acinetobacter baumannii* ATCC 19606 | ATCC |
| *Escherichia coli* C0002 | McMaster clinical isolate |
| *Serratia marcescens* C0016 | McMaster clinical isolate |
| *Proteus mirabilis* C0027 | McMaster clinical isolate |
| *Enterobacter cloacae* C0035 | McMaster clinical isolate |
| *Klebsiella oxytoca* C0036 | McMaster clinical isolate |
| *Klebsiella aerogenes* C0045 | McMaster clinical isolate |
| *Morganella morganii* C0052 | McMaster clinical isolate |
| *Citrobacter freundii* C0055 | McMaster clinical isolate |
| *Serratia odorifera* C0082 | McMaster clinical isolate |
| *Stenotrophomonas maltophilia* C0101 | McMaster clinical isolate |
| *Citrobacter amalonaticus* C0120 | McMaster clinical isolate |
| *Staphylococcus aureus* TCH1516 | ATCC |

| | |
|---|---|
| *Klebsiella pnuemoniae* ATCC 43816 | ATCC |
| *Klebsiella pneumoniae* MKP103 | (1) |
| *Escherichia coli* BW25113 | Laboratory strain |
| *Pseudomonas aeruginosa* MPAO1 | Wild-type strain, (2) |
| *Pseudomonas aeruginosa* MPAO1 PA4394::Tn5 | *mscS-1* mutant; Mutant strain identifier: phoAwp02q2D03, (2) |
| *Pseudomonas aeruginosa* MPAO1 PA4380::Tn5 | *colS* mutant; Mutant strain identifier: phoAwp02q4A03, (2) |
| **Plasmids** | |
| pUCP22 | (3); from J. Dennis lab, U. Alberta |
| pUCP22-MscS-1$^{WT}$ | This work |
| pUCP22-MscS-1$^{K73N}$ | This work |
| pUCP22-ColS$^{WT}$ | This work |
| pUCP22-ColS$^{T201P}$ | This work |
